# Regulation of the branched-chain amino acid pathway and its crosstalk to carbon metabolism in yeast

**DOI:** 10.1101/2025.05.05.652184

**Authors:** Ximena Escalera-Fanjul, Olufemi Bolaji, Bernhard Drotleff, Alexander DeLuna, Theodore Alexandrov, Edda Klipp

## Abstract

In *Saccharomyces cerevisiae,* the TORC1 pathway regulates the transition from rapid fermentative growth to respiration during the diauxic shift, by tightly coordinating energy- and biomass-producing pathways. Although leucine and other components of the branched-chain amino acid (BCAA) pathway are known TORC1 regulators, how the BCAA pathway is controlled across this transition and influences the crosstalk between central carbon and amino acid metabolism remains unclear. By integrating high-throughput flow cytometry and untargeted LC-MS metabolomics into a thermodynamically curated genome-scale model, we profiled protein levels, metabolite dynamics, and cellular context during the diauxic shift of a GFP-tagged strain library of the BCAA pathway in wild-type and mutants lacking non-essential genes for BCAA biosynthesis. This analysis revealed that the BCAA pathway operates in two branches, with only the leucine-committed branch exhibiting a fermentative signature aligned with TORC1 activity. We further identified key regulatory elements and showed that dysregulation of the BCAA pathway disrupts distant metabolic circuits, including central carbon metabolism. Our findings highlight the dynamic role of the BCAA pathway in metabolic network integration during the diauxic shift.

## INTRODUCTION

Cells modulate their metabolic internal state—biomass and energy production—to optimize their growth rate to the environmental conditions. Fermentation and respiration differ in their energy and biomass yield; funneling pyruvate into the TCA cycle can generate up to 15 times more ATP and yield five times more biomass per glucose molecule than fermentation^1^. However, fermentation produces more ATP per protein mass, meaning less investment to generate energy. Hence, an accelerated glycolytic flux, requiring a limited pool of enzymes is energetically inexpensive, leaving untapped energy for rapid growth during fermentative metabolism^2^. These contrasting strategies serve different purposes, e.g., competition with other microorganisms in the case of fermentation while respiration maximizes the amount of energy derived from a limited resource, both which are needed to ensure population success.

The ability of *Saccharomyces cerevisiae* to ferment sugars to ethanol under aerobic conditions—known as the Crabtree effect—depends on a high glycolytic flux to supply cells with energy^3^ and has been associated with a “make-accumulate-consume” strategy, that helps yeast outcompete other microorganisms in environments with limited resources^4^. Under such growth regime, the glucose catabolic repression response down regulates the tricarboxylic acid (TCA) cycle and oxidative phosphorylation^5^. Although the catabolic repression response decreases mass and energy fluxes through the TCA cycle and respiratory chain, the TCA cycle displays enough activity to provide the necessary precursors to sustain the rapid growth rates associated with this metabolic state, namely the exponential- or log-phase^6^. As glucose becomes limiting, cells undergo a so-called diauxic shift, a transient phase in which cells decrease their growth rate as they switch from fermentation to respiratory metabolism. During this transition, cells begin respiring the remaining glucose, and as the post diauxic shift progresses, the released ethanol is catabolized through the TCA cycle^3^. While transitioning to this new metabolic state in which TCA cycle provides cells with both energy and biomass, cells undergo a series of temporally organized changes. For instance, following the down regulation of glycolysis the TCA, glyoxilate cycle and oxidative phosphorylation pathways are activated.

One of the underling mechanisms driving the transition from log to stationary phase is mediated by the target of rapamycin complex 1 (TORC1) signaling pathway. During fermentative growth, TORC1 is active and inhibits the transition from log-phase to diauxic and stationary phase by, for example, activating PKA, which inhibits the catabolism of non-fermentable carbon sources such as ethanol through Snf1 negative regulation and by the phosphoinhibition of Rim15, which is involved in the establishment of stationary phase ^7,8,9^. The TOR pathway largely relies on the conserved Rag GTPases Gtr1/Gtr2 to perceive environmental cues, primarily by sensing the intracellular concentration of certain metabolites. In both yeast and mammals, leucine is a potent positive regulator of TORC1 and is sensed by this canonical mechanism. However, perturbing the leucine biosynthetic pathway—the branched chain amino acid (BCAA) pathway—specifically at Bat1 amino transferase, results in inactivation of TorC1 in a Gtr1/Gtr2-independent manner. Instead, this inactivation is linked to TCA cycle malfunction. This observation has led to the proposal that the BCAA pathway regulates both leucine and TCA cycle metabolism to control TORC1 signaling^10^.

Beyond its role in regulating TORC1, the BCAA has other features that renders it a good candidate to coordinate growth and energy metabolism. It integrates amino acid and carbon metabolism through shared intermediates. Its spatial organization links it to mitochondrial volume dynamics, which fluctuate during the fermento-respiratory shift, influencing reaction rates ^11-12^ . The pathway also exhibits complex regulation shaped by evolutionary gene duplications associated with *S. cerevisiae’s* evolution toward a fermentative lifestyle. Moreover, the metabolic flux through this pathway is finely tuned by multiple mechanisms, including feedback inhibition by individual amino acids (isoleucine, valine, and leucine), ATP-dependent modulation, and transcriptional control mediated by the α-IPM-responsive transcription factor Leu3^5,13 and 14^.

Despite considerable focus on the mechanisms by which BCAA pathway regulates TORC1 activity, it remains unclear whether the pathway responds to the diauxic shift, which regulatory elements are involved during this major metabolic transition, and how the BCAA pathway connects to other energy-producing or growth-related cellular pathways. Here, we combined experimental and computational methods to provide a comprehensive characterization of the pathway’s dynamics, detailing its response to the diauxic shift and the roles of its components in both intra-pathway regulation and broader cellular metabolism. We show that during growth progression the BCAA pathway is splitted into two branches, the Val/Ile branch displaying a fermento-respiratory profile, and the leucine-committed branch following a fermentative signature. While the auxotrophic deletion strains *ilv6*Δ*, bat2Δ* and *leu9Δ* showed no physiological effect; *bat1*Δ, *leu4*Δ and *oac1*Δ strains affected a distinct segment of the pathway and influenced other metabolic pathways, including amino acid biosynthesis and central carbon metabolism. We provide experimental and computational evidence that dysregulation of the leucine branch reroutes pyruvate toward the TCA cycle, resulting in increased flux through pyruvate dehydrogenase (PDH) as well as the TCA and glyoxylate cycles during logarithmic growth. These findings indicate that the leucine-committed branch of BCAA metabolism regulates respiratory activity through its enzymatic function. In contrast, in the respiratory conditions, the metabolic response involves additional regulatory mechanisms beyond enzyme activity, suggesting the involvement of higher-level processes such as transcriptional regulation.

## RESULTS

### The BCAA pathway

Several features place leucine at the interface between biomass and energy metabolism. The BCAA pathway comprises 12 enzymes that convert threonine or pyruvate into isoleucine, valine or leucine^14^ (Fig. 1A). This pathway is well connected to central carbon metabolism, as three intermediates—pyruvate, ac-CoA and α-ketoglutarate (KG)—are shared between the two. Additionally, the pathway is compartmentalized in the mitochondria and the cytoplasm. The upstream portion, which converts threonine or pyruvate to α-isopropylmalate (α-IPM), valine or isoleucine, is confined to the mitochondria, while the downstream segment, converting α-IPM to leucine occurs in the cytosol. Mitochondrial morphology and volume change according to the metabolic state of the cell^15^ and the rates of the chemical reactions embedded in this compartment are directly affected by its volume ^11^ ^and^ ^12^. Furthermore, two pairs of paralogous proteins—Bat1/Bat2 and Leu4/Leu9—that arose from the allopolyploidization event that facilitated *S. cerevisiae* fermentative lifestyle^5,13^ ^and^ ^14^, are involved in branching points of the pathway and are associated with negative feedback regulation.

**Figure 1.**
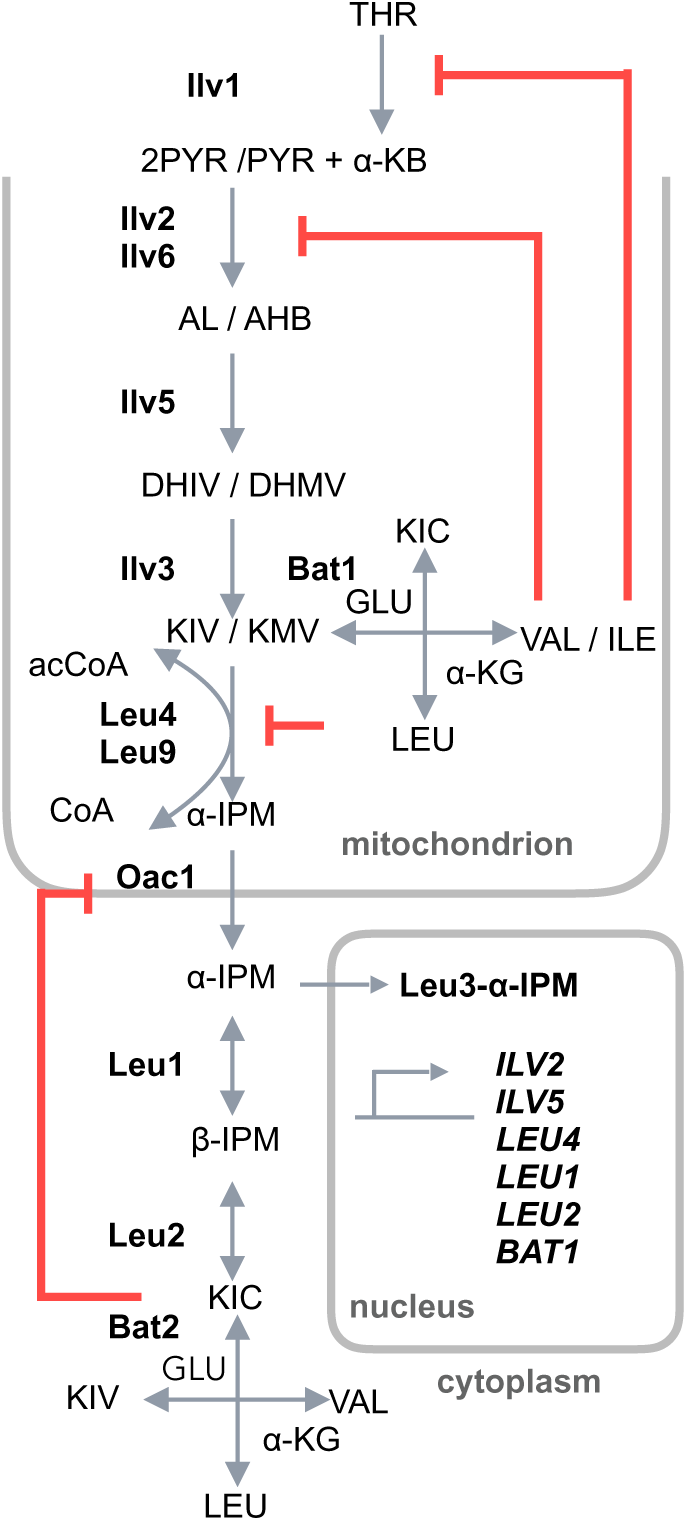
The BCAA pathway and its regulation. Arrows indicate irreversible (simple) and reversible (double) reactions. Enzymes (bold), metabolites (uppercase), and Leu3–α-IPM-activated genes (uppercase italics). Enzymes: Ilv1 (threonine deaminase), Ilv2/ Ilv6 (acetohydroxy acid synthase catalytic/regulatory subunit), Ilv3 (dihydroxy acid dehydratase), Bat1/Bat2 (branched-chain amino acid aminotransferase mitochondrial/cytosolic); Leu4/Leu9 (α-isopropylmalate synthase), Leu1 (isopropylmalate isomerase), Leu2 (β-isopropylmalate dehydrogenase). Abbreviations: THR (threonine), PYR (pyruvate), α-KB (α-ketobutyrate), AL (acetolactate), AHB (2-aceto-2-hydroxybutyrate), DHIV (dihydroxyisovalerate), DHMV (β-dihydroxy-β-methylvalerate), KIV (α-ketoisovalerate), KMV (α-ketomethylvalerate), VAL (valine), ILE (isoleucine) α-IPM (α-isopropylmalate), β-IPM (β-isopropylmalate), KIC (α-ketoisocaproate), aCoA (acetyl coenzyme A) CoA (coenzyme A), αKG (α-ketoglutarate) and GLU - glutamic acid.

The BCAA pathway system has five major regulatory features that likely fine-tune metabolic flux through the pathway: i) Selective isoleucine feedback inhibition: The first enzyme in the pathway (Ilv1) is inhibited by Isoleucine ^16^ ii) Selective valine feedback inhibition: The second enzyme in the pathway consists of the catalytic and the regulatory subunits (Ilv2 and Ilv6, respectively). While the catalytic subunit alone is active and insensitive to valine inhibition, its association with the regulatory subunit stimulates its activity, but renders it sensitive to valine inhibition. This inhibition can be reversed by ATP^17^. iii) Distinct leucine feedback inhibition: The first committed step for leucine production is catalyzed by the Leu4 and Leu9 paralogs, which can form homodimers or, preferentially, heterodimers. These three possible isoforms differ in their sensitivity to leucine: the Leu4-homodimer is leucine-sensitive, the Leu9-homodimer is leucine-resistant, and the Leu4-Leu9 heterodimer has intermediate sensitivity^13^. iv) α-IPM transport inhibition: The mitochondrial transporter Oac1, responsible of exporting α-IPM, is inhibited by α-ketoisocaproate (KIC), the immediate precursor of leucine ^18^ . v) α-IPM transcriptional regulation: The transcription factor Leu3 regulates the expression of several genes in the Leu/Val pathway. Leu3 acts a dual regulator, which in the presence of α-IPM acts as transcriptional activator and, in its absence, acts as repressor ^14^ ^and^ ^19^.

### High-throughput measurement of the BCAA pathway protein profile in wild type and knockout strains

To analyze how the protein levels of the BCAA pathway adjust to the fermento-respiratory shift and to genetic perturbations, the fluorescent intensity of a library of GFP-fusions, in wild-type and knockout background strains, was followed by high-throughput cytometry across different growth phases (from early log-phase to post diauxic shift; Fig. S1). The library was generated by mating an array of Ilv2-, Ilv6-, Ilv3-, Bat1-, Bat2-, Leu9-, Oac1-, Leu1-, Leu2- and Rpl41b-GFP fusion strains (Fig. 2A), to a query collection composed of all deletion strains of the BCAA pathway that were not associated with a BCAA auxotrophy *ilv6*Δ*, bat1*Δ*, bat2Δ, leu4*Δ*, leu9Δ* and *oac1*Δ (Fig. 2B).

**Figure 2.**
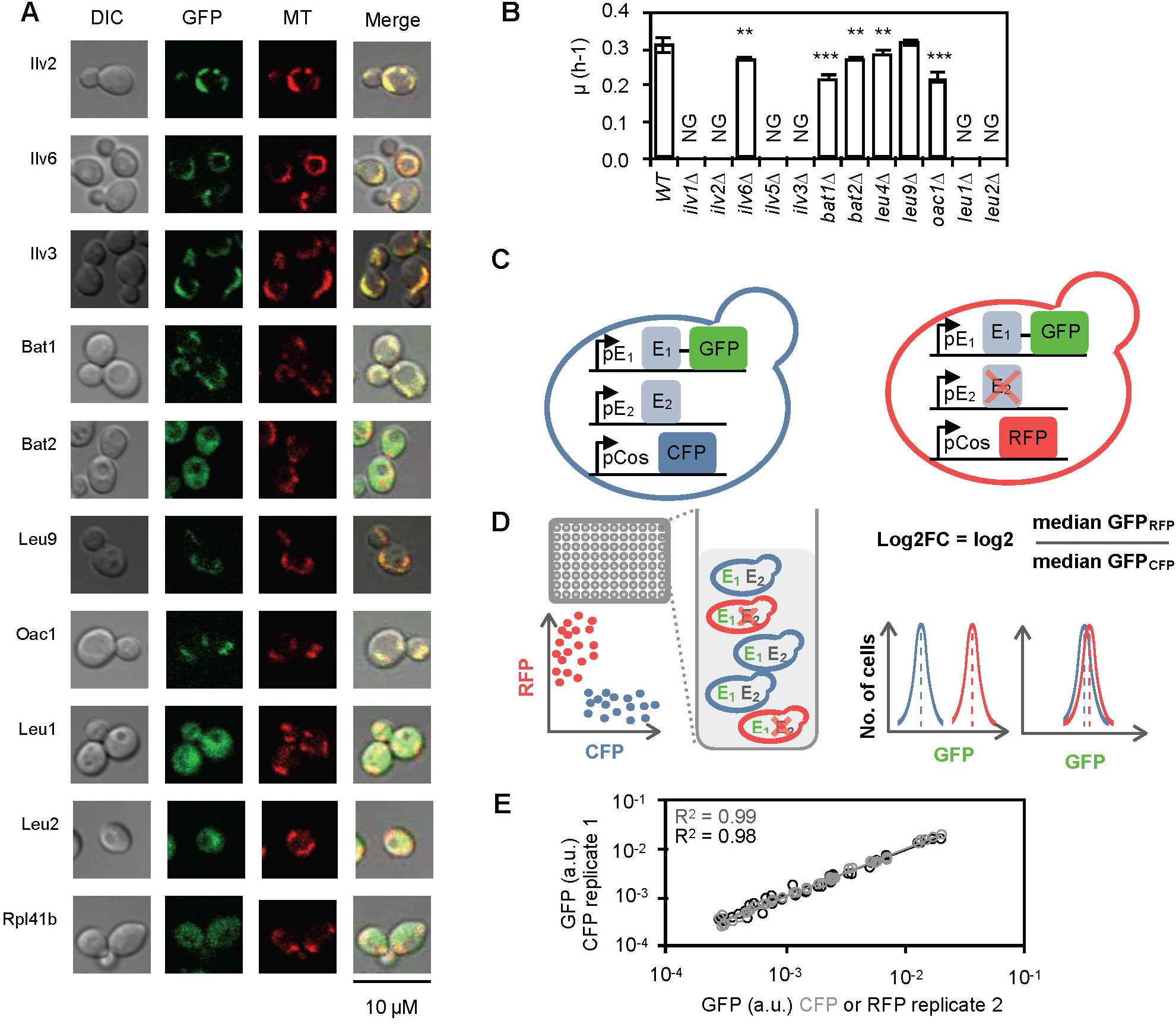
Experimental setup to assess the protein profile of the BCAA pathway. **A)** Cellular localization of GFP fusions strains grown in SD media to exponential phase (OD600 of ∼0.6). Mitochondrial staining with MitoTracker CMXRos to determine sub cellular localization by confocal microscopy. DIC, differential interference contrast. Representative image of multiple cells from two biological replicates. **B)** Growth rate analysis of wild type and deletion strains, values represent means ± SD from three independent experiments. NG: no growth. Asterisks indicate significant growth differences from wild-type (one-tailed, homoscedastic t-test: ∗*p* < 0.01; ∗∗*p* < 0.001; ∗∗∗*p* < 0.01). **C)** Schematic of the experimental strategy: each BCAA enzyme (E₁→ₙ) was GFP-tagged in either a wild-type (constitutively expressing CFP) or a deletion strain (expressing RFP). **D)** Wild-type and deletion strains were co-cultured in 96-well plates and analyzed by three-color flow cytometry to distinguish strains and measure GFP intensity. Median GFP levels served as a proxy for protein abundance and were used to estimate Log₂ fold changes (Log₂FC) of the median GFP intensity ratio of deletion to wild-type strains present in the co-culture. **E)** Dye-swap control showing median GFP intensity from two wild-type biological replicates over the complete time series, independently expressing CFP (gray) or independently expressing CFP or RFP (black). R^2^: Pearson’s linear correlation coefficient.

To ensure the growth of the wild type and knockout strains under identical environmental conditions, the fluorescent protein cerulean (CFP) was constitutively expressed in the wild-type strains and the mCherry (RFP) in the knockouts (Fig. 2C). The distinctive fluorophores allowed us to identify by high-throughput flow cytometry wild type and knockout cells in a co-culture, while following the GFP signal in each cell type (Fig. 2D). The GFP signal of each cell was normalized by its relative cell size (FSCA) and the median GFP intensity of the population was used as a proxy of the protein abundance (Fig. 2D).

To assess whether the different fluorophore backgrounds affected the GFP levels, we included a set of wild-type strains in a RFP background. A non-parametric statistic (Wilcoxon test, p < 0.01, Supp Data 1-sheet CFPwt-RFPwt -WT) failed to identify any significant differences in the protein abundance between wild-type strains expressing CFP or RFP co-cultured in synthetic minimal medium. Accordingly, the wild-type protein abundances between independent replicates in CFP or RFP backgrounds showed a high correlation throughout the different growth phases (Fig. 2E).

### The BCAA pathway operates two branches with contrasting responses to diauxic shift

As mentioned above, we followed the fluorescence intensity at different time points tracking changes in protein levels across the growth phases, from the log phase to the post-diauxic shift. This time-resolved approach allowed us to capture the protein abundance dynamics of the BCAA pathway during the fermento-respiratory transition. In the wild-type strain the most abundant enzymes of the BCAA pathway were Ilv3 and Leu1, with median GFP intensities as high as 0.07 and 0.16 a.u., respectively. In contrast, Bat2 and Leu9 were the enzymes with the lowest abundances, displaying minimum median GFP intensities around 0.003 a.u. The remaining pathway components Ilv2, Ilv6, Bat1 and Oac1 were present at intermediate levels, ranging from 0.01 - 0.03 GFP a.u. (Fig. 3A and Table S1). The rate between the maximum and minimum values of the median protein abundances across the time series identified Ilv2, Leu1 and the reference protein Rpl41b as the enzymes that displayed the least pronounced responses to the growth phase progression (Fig. 3A and Table S1). On the contrary, Oac1 and Ilv3 displayed the most prominent responses, halving and triplicating their abundance, respectively, while the growth phases progressed. The remaining enzymes Ilv6, Bat1 and Bat2 increased around 50% and Leu9 decreased approximately 25% its abundance throughout the time course tested (Fig. 3A and Table S1).

**Figure 3.**
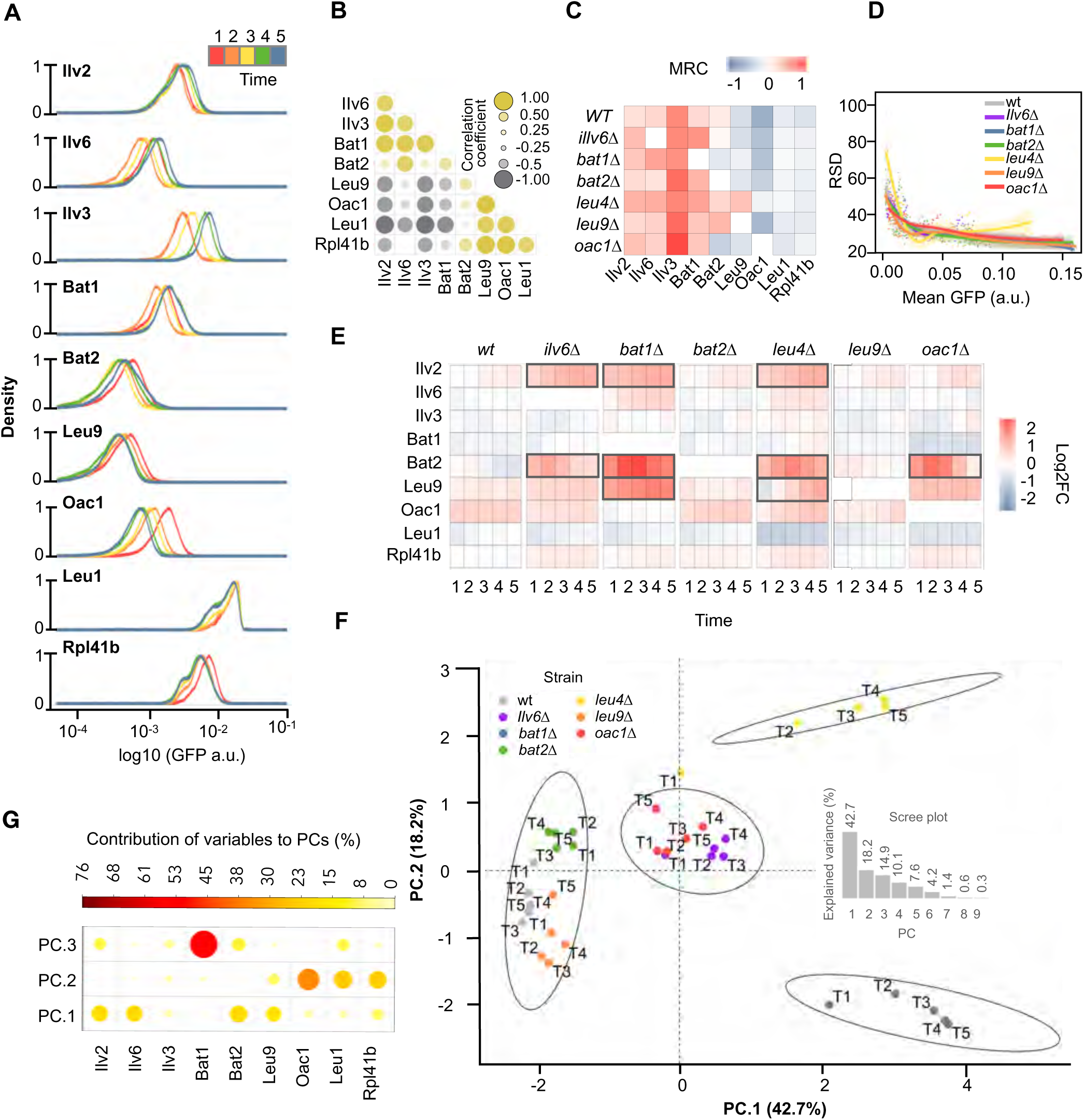
Time course of the BCAA pathway protein profile in wild-type and knockout strains. A) Wild-type distribution of the protein levels of proteins of the pathway. Protein abundance density plot obtained by flow cytometry of a replicate of the wild-type strain expressing RFP constitutively. **B)** Crosscorrelation analysis in the wild type strain. Values estimated from the mean protein levels across the time series of three independent experiments **C)** Mean relative change of the protein abundance was estimated as the average of the differences between the median GFP intensity of successive time points of three independent experiments. **D)** Loess curve of the mean coefficient of variation of wild-type and deletion strains considering each biological replicate independently and the complete time course. Mean (solid lines) and standard error (shaded area) of three biological replicates. **E)** Heatmap capturing the dynamics of the protein profile response in deletion strains. Log2FC from the mean of three independent biological replicates. Boxes highlight components significantly affected (MB-test p < 0.01 and cutoff of doubling or halving the abundance). **F)** PCA of wild-type and knockouts across growth phases shows deletions impacting pathway profiles. on the first two eigenvectors of PCA (PC1, PC2) and k-means clustering. **G)** PCA loadings representing proteins that drove the segregation between the clusters.

A cross-correlation analysis was used to quantify the similarity between protein abundance temporal patterns. The analysis revealed that the enzymes involved in the biosynthesis of the three amino acids (Ilv2, Ilv6, Ilv3, Bat1 and Bat2) displayed a similar temporal profile (Fig. 3B). An initial drop in their abundance, followed by a consistent increase as the time course progressed. Their abundance peaked at the final two time points, which correspond to the diauxic and post-diauxic phases (Fig. 3A). We named this branch of the pathway the “shared-branch. In contrast, the proteins committed exclusively to leucine biosynthesis (Leu9, Oac1 and Leu1) showed a decrease in concentration throughout the growth phases (Fig. 3A and S4). Even Leu1, which followed a bimodal distribution, favored the lower abundance peak in the later time points (Fig. 3A). The cross-correlation values within each group were high, indicating similar patterns over time (Fig. 3B). Although the reference protein Rpl41b displayed rather small abundance changes, its abundance decrease resulted in high correlation values with the leucine committed proteins (Fig. 3B). The negative cross-correlation values between the two groups, i.e., the shared- and the leucine committed-proteins, indicated opposing profiles, i.e., while one branch increased the other decreased (Fig. 3 A-B). Relevantly, the contrasting profiles, between the two groups of proteins, were not related to the sub-cellular localization of the enzymes (Fig. 1A and 2B).

To assess whether the growth phase signature of the pathway was robust to genetic perturbations, we estimated, for each protein in the pathway, the mean relative change between successive time points in both the wild-type and knockout strains. Although the initial abrupt decrease in the concentration of the shared enzymes — likely marking the end of the lag phase — partially masked their overall increasing trend (Fig. 3A), calculating the mean relative change effectively compressed the time-series data. This simplification enabled us to better evaluate the robustness of the BCAA pathway’s growth phase signature in response to genetic perturbations (Fig. 3C). Like the wild type, most of the knockout strains displayed the same contrasting profile, in which the concentration of the shared enzymes was up-regulated and the abundance of leucine committed proteins were down-regulated while the growth phases progressed. Only few elements reversed their abundance profile in a consistent manner and distanced from the wild type behavior, i.e., Bat2 upon *bat1Δ* and *oac1Δ* deletions, as well as Leu9 in the *leu4Δ* background (Fig. 3C and S3).

Next, we asked how robust the system was in general to genetic perturbations. Testing the relative standard deviation (RSD) of all the proteins of the pathway at all time points, revealed that the mean abundance of the proteins of the pathway anti-correlated with their level of dispersion. A steep increase in the RSD of protein levels was observed as protein abundance decreased, continuing until a threshold was reached. Beyond this threshold, further reductions in protein abundance did not result in a corresponding increase in RSD, effectively defining a lower bound for protein noise (Fig. 3D and Supp Data 1) (Jakub Jędrak* & Anna Ochab-Marcinek, 2020). However, in the *leu4Δ* deletion strain we observed a general increase in the coefficient of variation, particularly in the more concentrated enzyme Leu1. In this knockout background, a subpopulation of cells decreased the concentration of Leu1, the shift towards lower levels favored the low intensity peak, and such a displacement resulted in a platykurtic distribution with high RSD (Fig. 3D, S3 and Supp Data 1).

### Genetic perturbations over the BCAA pathway generate distinctive protein signatures

The temporal dynamics of the protein profile, showing the direction and intensity of the changes in the knockout strains (Fig. 3E), were used to identify, which responses were significantly affected (Supp Data). After performing a normality test over the bootstrapped log2-fold change (log2FC) mean distribution (Shapiro-Wilk, p > 0.05, Supp Data), a combined multivariate statistic MB-statistic^20^, designed to compare temporal series (F-test based on multivariate empirical Bayes statistic, adjusted p-value < 0.01), and a cutoff of doubling or halving the protein concentration, identified 9 out of the 30 responses as significantly different from the wild type.

While *bat2Δ* and *leu9Δ* protein profiles did not exhibit significant differences compared to the wild type, *ilv6Δ*, *bat1Δ*, *leu4Δ* and *oac1Δ* knockouts triggered significant responses in other components of the pathway. It is worth noting that the increase in RSD as protein levels drop may obscure the significance of downregulated elements, such as Oac1 and Leu1, following genetic perturbation (Fig. 3D). Having this in mind, the proteins that were significantly affected were Ilv2, Bat2, and Leu9, that happened to be mostly up-regulated. The remaining proteins, including the reference protein Rpl41b, did not respond in a significative manner to any of the genetic perturbations tested. Interestingly, each knockout affected a different set of proteins and with different strength depending on the growth phase, resulting in a unique protein profile signature for each knockout (Fig. 3E). Ilv2 exhibited a cumulative response, with the highest difference to the wild type strain at later time points. Bat1 response was more intense at time point 2 and 3, corresponding to the late log phase. In contrast, Leu9 displayed varied responses to different deletions. While it showed a general up-regulation in response to *bat1Δ* deletion, it exhibited an initial down-regulation followed by a consistent up-regulation in the *leu4Δ* deletion background (Fig. 3E).

In order to analyze the overall response of the protein profile and not protein by protein, we performed a principal component analysis (PCA, Fig. 3F). The projection of the log2FC time course onto the first two PCA eigenvectors revealed that knockout strains appear in four defined clusters, identified by k-means clustering method. The first cluster was composed by the knockouts *bat2Δ* and *leu9Δ* together with the wild-type, implying that in these deletion backgrounds there were only small changes in their protein profiles. A second cluster with low correlation values to the first and second principal components (PC1 and PC2, respectively) included *ilv6Δ* and *oac1Δ* knockout strains. *ilv6Δ* and *oac1Δ* strains were distinguished from each other by the third principal component (PC3; Fig. 3F and S4). The third and the fourth clusters were respectively comprised of *bat1Δ* and *leu4Δ* and were only distinguished form each other by the PC2. PC1 captured 43% of the total variance and was able to distinguish three out of the four cluster (clusters 1, 2 and 3-4, Fig. 3F and inset),. Ilv2, Ilv6, Bat2, and Leu9 were the enzymes that influenced this cluster most (Fig. 3G). The transporter Oac1 contributed the most to PC2, whereas Bat1 was the element that mainly influenced PC3 (Fig. 3G). Together, the first three components accounted for 75% of the variance (Fig. 3F inset) and were able to distinguish the knockout strains populations in agreement with the MB-statistic, supporting that the metabolic changes induced by *ilv6Δ*, *bat1Δ*, *leu4Δ* and *oac1Δ* knockout strains were different.

### The BCAA Pathway Operates in a Branched Manner with Distinct Coupling to Central Carbon Metabolism: Leucine Branch fermentative vs Valine/Isoleucine Fermento-Respiratory

To assess the physiological changes at metabolic level, we analyzed the metabolic profiles of the wild-type and knockout strains, from log-phase to post-diauxic shift, using an untargeted metabolomic approach based on LC-MS in positive and negative mode. Metabolites present in both modes displayed highly similar profiles (Fig. S5) and reproducibility of replicate measurements was high, Spearman’s rank correlation coefficient always above 0.75 (Table S2 and Fig. S6-7). We detected 127 different metabolites (74 and 53 in positive and negative mode, respectively) that fell into various KEGG categories: energy, lipid, cofactors and vitamins, nucleotides, carbohydrates and amino acids. Metabolites related to secondary metabolism and orphan metabolites were classified as “others” (Fig. 4A).

**Figure 4.**
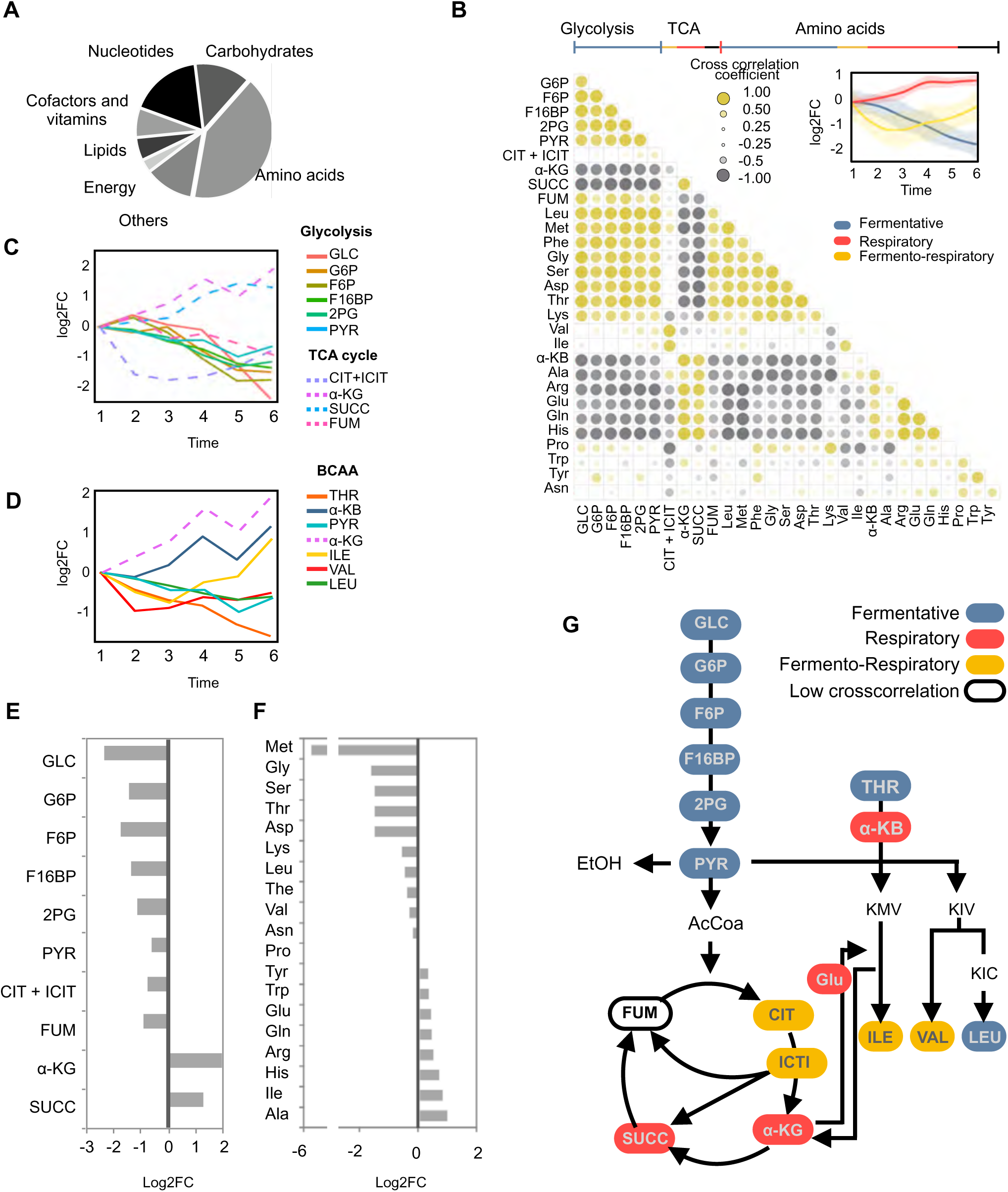
Wild type metabolic dynamiscs over diauxic shift. **A)** Untargeted LC-MS metabolomics detected 127 metabolites (74 in positive and 53 in negative mode) mapped to KEGG pathway categories; amino acid and unclassified metabolites were grouped as “others.” **B)** Cross-correlation analysis showed that 69% of metabolites followed similar temporal profiles, grouped into fermentative, fermento-respiratory, or respiratory categories (inset: loess curves). Categories were defined by central carbon metabolite clusters with correlation > 0.75; other metabolites were assigned to a category if their average correlation with that group exceeded 0.75 Full metabolite set in Supp Data 3). Loess curves show the mean ± SEM of carbon and amino acid metabolites. **C -D)** Loess curve showing the detailed temporal profile of from three biological replicates. **E and F)** Log₂ fold change of the intensity of metabolite concentration between the first and last time points from each metabolite’s loess fit. **G)** Schematic representation of the metabolic response of the BCAA pathway and central carbon metabolism based on the cross correlation analysis.

Next, to identify metabolites synchronized through the diauxic shift, we calculated correlations between metabolite levels over six timepoints. This cross-correlation analysis identified that 70% of the detected metabolites profiles fall into three specific categories that agree with central carbon metabolism temporal behavior:, i.e., fermentative, respiratory, and fermento-respiratory (Fig. 4B, 3B-inset). Fermentative metabolites comprised glycolytic metabolites and several amino acids, including leucine and methionine (Fig. 4B). The high cross-correlation coefficient values within this group resulted from their temporal signature, i.e., a constant concentration decrease as the growth passes progressed (Fig. 4B-inset, C-D). Respiratory metabolites included α-ketoglutarate and succinate, which are both lower TCA cycle metabolites, and other amino acids including glutamic acid and alanine (Fig. 4B). The temporal signature of this group was a constant increase as the growth phases progressed reaching a plateau at later growth phases (Fig. 4B-inset, C). These canonical fermentative and respiratory behaviors resulted in a strong negative cross-correlation between these two groups (Fig. 4B). The third group, here referred to as fermento-respiratory, was composed of cis-aconitate, valine and isoleucine (Fig. 4B). The temporal signature of this small group of metabolites was characterized by an initial decline in concentration followed by a constant increase while the growth phases progressed (Fig. 4B-inset, C-D), suggesting a hybrid behavior, initially fermentative and later respiratory. The full metabolite list in each category can be found in Supp Data 3.

In general, the metabolites belonging to upper glycolysis displayed stronger fermentative signatures than lower glycolytic metabolites, with glucose decreasing to a quarter of its initial concentration between log-phase and post-diauxic shift (Fig. 4E). Instead, metabolites belonging to lower glycolysis as well as upper TCA cycle metabolites exhibited weaker patterns (i.e. pyruvate and fumarate), or even hybrid signature such as cis-aconitate (Fig. 4C and F). In contrast, lower TCA cycle metabolites followed a canonical respiratory concentration increase, approximately doubling their concentration as the growth phases progressed (Fig. 4F). The amino acids that exhibited the strongest fermentative patterns were Gly, Ser, Thr and Asp (note that methionine was externally supplemented). The strongest respiratory metabolite was alanine (Fig. 4F), in the case of valine and isoleucine, that exhibited fermento-respiratory hybrid patterns, valine has demonstrated more fermentative behavior, whereas isoleucine demonstrated a shorter but stronger respiratory signature (Figs 4D and 4F). It is worth mentioning that the growth phase dependent profile of leucine concentration (fermentative) and that of valine and isoleucine (fermento-respiratory) matched the protein profile of the Leu- and the shared-branch of the pathway, respectively (Fig. 3A). This represents manifestation of two different behaviors for the two branches of the BCAA pathway.

### Genetic perturbation of the BCAA pathway affects other metabolic pathways, including TCA- and glyoxylate-cycle

To explore the impact of the knockouts on the metabolic profile of BCAA pathway with temporal resolution, we employed the MB-statistic^20^ (F-test based on multivariate empirical Bayes statistic, adjusted p value < 0.05). *bat2Δ* and *leu9Δ* knockouts did not produce significant alterations in the metabolic profile of the BCAA pathway, consistent with their protein profile signature (Fig. 3E). Despite the fact that *ilv6Δ* deletion led to changes in the protein profile, these alterations were not reflected at the metabolic level. This suggests that the protein changes upon *ilv6Δ* deletion can be compensated for any metabolic imbalance. In contrast, the protein profiles observed after *bat1Δ*, *leu4Δ*, and *oac1Δ* deletions (Fig. 3E) aligned with the metabolic changes (Figs 5A). The deletion of *bat1Δ* led to significantly decreased levels of valine and isoleucine and to leucine accumulation. In fact, the temporal patterns of valine and isoleucine were similar and mirrored that of leucine. In the *leu4Δ* knockout strain, lacking one of the two enzymes required in the first committed step of leucine biosynthesis, we observed sustained reduced levels of leucine across all the growth phases compared to the wild type. Lastly, *oac1Δ* deletion exhibited a significant reduction in threonine and leucine concentrations (Fig. 5A). Upon examining the three deletion strains — *bat1Δ*, *leu4Δ*, and *oac1Δ* — and their respective responses, it became evident that the pathway responded to each genetic perturbation in a specific manner at both the proteomic and metabolic levels.

**Figure 5.**
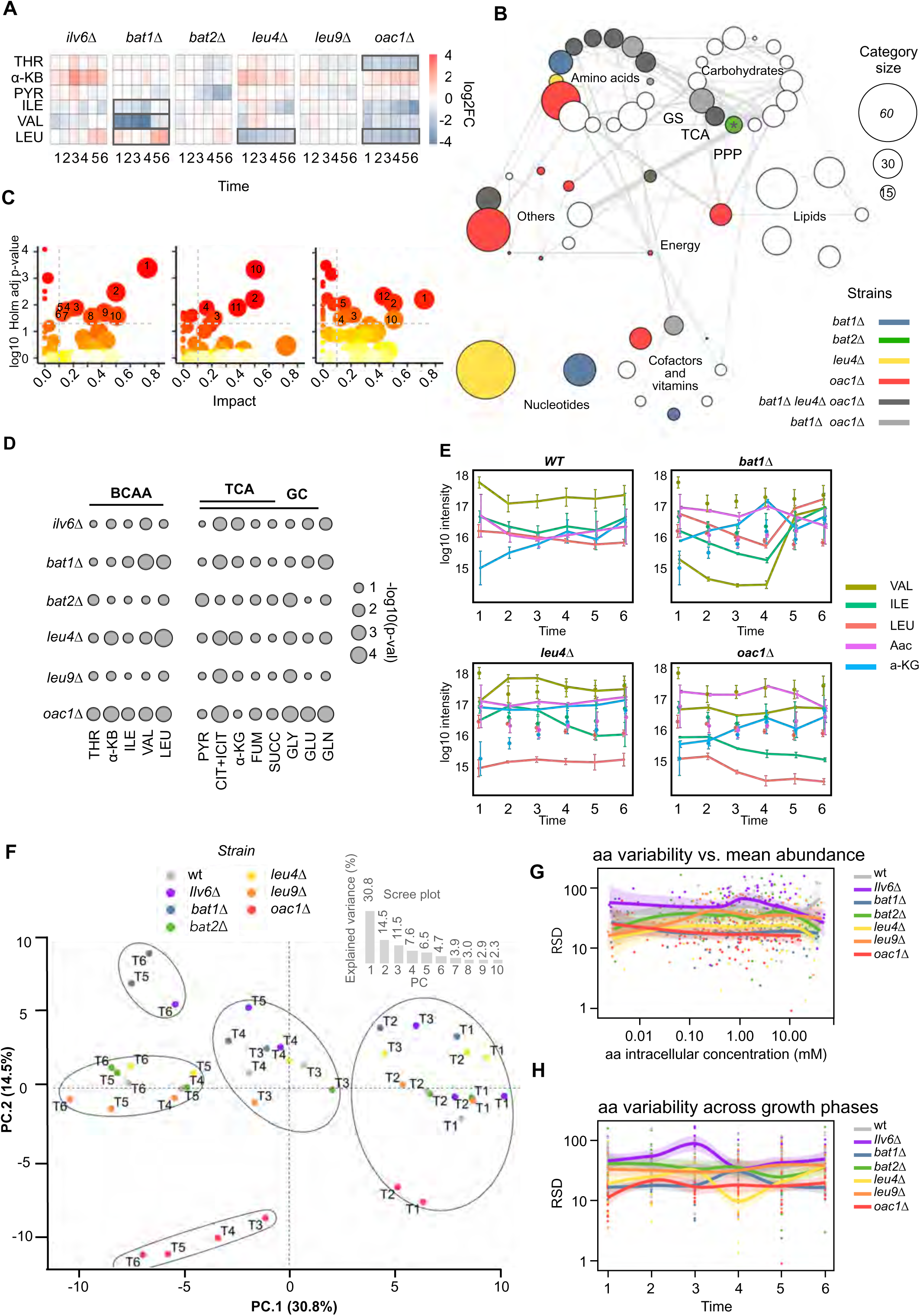
Genetic perturbations impact metabolism beyond the BCAA pathway. **A)** Heatmap capturing the dynamics of the BCAA pathway metabolic profile in deletion strains. Log2FC from the mean of three independent biological replicates. Boxes highlight components significantly affected (MB-test p < 0.01 and cutoff of doubling or halving the abundance). **B)** Metabolite enrichment analysis of metabolites at the time point with strongest Log2FC relative to wild type. Nodes: KEGG metabolite modules of measured metabolites, node size: metabolite count, edge weight: count of shared compounds, colored nodes: significant perturbations (Holm-adjusted p < 0.05): asterisks: false positives based on ranked list inspection and orthogonal validation. GS (glyoxilate cycle) TCA (Tricarboxiclic acid cycle) and PPP (pentose phosphate pathway). **C)** Pathway analysis based on the Holm statistic associated adjusted p-value < 0.05 and metabolite pathway impact < 0.1. Significant pathways are 1. Alanine, aspartate and glutamate metabolism, 2. Arginine biosynthesis, 3. TCA cycle, 4. Valine, leucine and isoleucine biosynthesis 5. Glyoxilate and decarboxylate metabolism 6. Biotin, 7. Pantothenate and CoA biosynthesis, 8. Arginine and proline metabolism, 9. Pyrimidine metabolism, 10. Glutathione metabolism, 11. Purine metabolism and 12. Cysteine and methionine metabolism. **D)** Metabolite significance based on p-value. **E)** Time-course of the mean values +/- SD of relevant metabolites. Wild-type values shown in knockout panels as points for reference. **F)** PCA of wild-type and knockouts across growth phases shows deletions impacting pathway profiles. on the first two eigenvectors of PCA (PC1, PC2) and k-means clustering. **G -H)**. Loess curve of the mean coefficient of variation of wild-type and deletion strains considering each biological replicate independently and the complete time course. Mean (solid lines) and standard deviation (shaded area) of three biological replicates.

We expanded our analysis to explore to which extent the genetic perturbation affected other parts of the metabolism. The MB-statistic^20^ (F-test based on multivariate empirical Bayes statistic, adjusted p value < 0.05) identified that *bat1Δ*, *leu4Δ* and *oac1Δ* had significant impact beyond the BCAA pathway, including other amino acids and metabolites associated to carbohydrate metabolism. Each knockout significantly affected 44 (*bat1Δ*), 24 (*leu4Δ*), and 43 (*oac1Δ*) out of the 127 detected metabolites (Supp Data 3). To relate the affected metabolites to a biologically relevant context, we conducted an enrichment analysis considering the metabolite intensities of the time point with the strongest log2FC relative to the wild type (Fig. 5B). In line with the MB-statistic, the enrichment analysis (Holm statistic adjusted p-value < 0.05) identified that *bat1Δ*, *leu4Δ* and *oac1Δ* had broad metabolic effects, influencing pathways outside of the BCAA pathway. The three strains widely overlap in the pathways that were affected within both amino acid and carbohydrate metabolism, i.e. cysteine and methionine, arginine and proline, phenylalanine, tyrosine and tryptophan biosynthetic pathways as well as TCA cycle. *bat1Δ* and *oac1Δ* further overlapped in alanine, asparagine and glutamic acid metabolism, as well as glyoxylate and dicarboxylate metabolism. While *leu4Δ* affected some pathways in a specific manner it was clear that its effect was more restricted than upon *bat1Δ* deletion or especially in comparison to the *oac1Δ* strain, which affected several metabolic pathways beyond amino acid and carbohydrate metabolism (Fig. 5B and S8). Unlike the longitudinal MB-statistic, the cross-sectional approach falsely identified an effect of the *bat2Δ* deletion on the pentose phosphate pathway. Closer inspection showed this was a false positive, caused by the built-in arbitrary *p*-value cutoff of the ranked list of metabolites. This highlights the strength of longitudinal analysis and the value of using complementary analytical methods

A topology-based pathway analysis highlights metabolic pathways significantly disregulated by changes in central metabolites — those occupying key positions and exerting major influence within the pathway network. (Fig. 5C, MetaboAnalyst 3.0 Impact analysis Holm adjusted p-value < 0.05 and impact > 0.1). Arginine biosynthetic pathway and alanine, asparagine and glutamic acid metabolism had several significantly affected metabolites in central positions (e.g. N-Acetylornithine, L-Citrulline and L-Ornithine in arginine biosynthetic pathway and N-Carbamoyl-L-aspartate in the case of alanine, asparagine and glutamic acid metabolism). Although these pathways shared around 50% of their detected metabolites, they were affected in different elements in each of the knockouts, and the affected elements in each of the deletion strains did not completely overlap in these two pathways (Supp Data 3). This finding emphasizes the fact, that the specificity of the response to each perturbation was extended to other parts of the metabolism. In cases where two deletion strains affected the same pathway, they still altered different components within it— underscoring the unique metabolic impact of each mutation.

Other pathways, such as BCAA and TCA cycle, showed high impact values, albeit lower than the two pathways described above, due to the mid-low importance of the detected metabolites within these pathways, as defined by the topology outlined in KEGG (Fig. 5C). A detailed examination of the BCAA biosynthetic pathway revealed that *bat1Δ* preferentially affected the shared-branch, *leu4Δ* the Leu-branch and *oac1Δ* the overall pathway. In the TCA cycle, *bat1Δ* and *leu4Δ* not only impacted the pathway at the level of α-ketoglutarate but also influenced cis-aconitate. In contrast, *oac1Δ* only affected the latter. It is worth mentioning that the effect over the glyoxylate cycle seemed to be determined by the overlap it has with the TCA cycle metabolites, in addition to some amino acids (Fig. 5D). This highlights the potential impact that perturbations on the BCAA pathway could have over the fluxes of a peroxisomal metabolic pathway ^21^.

The detailed analysis of metabolite dynamics within the BCAA and TCA pathways revealed that in the *bat1Δ* strain isoleucine and valine concentrations decreased during logarithmic growth, while cis-aconitic acid and α-ketoglutarate concentrations increased (Fig. 5E upper left). However, while the cell culture transitioned to diauxic and post-diauxic shift, the concentration of these metabolites returned to wild-type levels, coinciding with an increase in leucine concentration. This observation suggest, that the *bat1Δ* mutation resulted in a flow block that clogged TCA-cycle flow around the point of Aco1 and α-ketoglutarate dehydrogenase (α-KGDH) action, which was later relieved by the up-regulation of the Leu-branch, which drained the accumulated TCA-cycle metabolites. Conversely, in the *leu4Δ* deletion strain, leucine concentration was consistently lower than in the wild type, and cis-aconitate and α-ketoglutarate were accumulated through all growth phases, potentially due to the lack of up-regulation of the shared-branch and the incomplete compensation of the Leu9-branch (Fig. 3E, 5A and E). Finally, in the *oac1Δ* knockout, the decrease in BCAA concentration became more severe as the growth phases progressed, while cis-aconitate accumulation was gradually alleviated (Fig. 5E). Together, these findings show that beyond the pathway- or element-specific specific effects triggered by each genetic perturbation, the impact of these perturbations can also vary in intensity across different growth phases.

In agreement with these observations, a PCA analysis (Fig. 5F) revealed that at diauxic - post diauxic phase (timepoints T5 and T6) the strains *bat1Δ* and *oac1Δ* were more different from the wild type than at log phase (timepoints T1-T4). The projection of the metabolite intensities onto the first two PCA eigenvectors revealed that the PC1 was able to resolve the different growth phases for all strains, while PC2 distinguished *oac1Δ* knockout from the rest of the strains, as well as *bat1Δ* and surprisingly *ilv6Δ* at diauxic-postdiauxic phase. These two PCs explained 45.3% of the variation (Fig. 5F-inset) and were able to capture that, - while the growth phases progressed, - the difference driving PC2 was more pronounced for *oac1Δ* and *bat1Δ* strains than for the remaining strains. The contribution to PC1 was distributed among several metabolites from different metabolic pathways, encompassing several glycolytic elements and amino acids with a glycolytic profile (Fig. S9). A mosaic of metabolites falling into different KEGG categories, including respiratory amino acids and non-proteinogenic amino acids contributed the most to PC2 (Fig. S9), suggesting that the growth phase specific response observed in *bat1Δ* and *oac1Δ* was extended to other metabolic pathways. Furthermore, the extended yet mild response to *ilv6Δ* deletion at later time points, rendered challenging for detection using stringent methods, but was captured by the PCA. On the contrary, *leu4Δ* that showed a more limited response than that of *bat1Δ* and *oac1Δ*, was not distinguishable from the wild type by this analytical method.

The use of internal standards allowed us to quantify the intracellular concentrations of 15 amino acids in the wild type strain. We used these data to interpolate the amino acid intracellular concentrations in a semi-quantitative manner for the rest of the strains (Materials and Methods). In the wild type strain, the most concentrated amino acids regardless of the growth phase were alanine and glutamate, with the maximum mean concentrations of 40 mM and 27 mM, respectively. This was three orders of magnitude higher than the lowest concentrated amino acid methionine at 0.03 mM (Supp Data 3). Estimated concentrations were in agreement with those reported by (Mülleder, 2016, Pearson correlation coefficient of 0.93). Consistent with Mülleder 20016, but in contrast to what was observed with the protein concentration, we did not observe a correlation between amino acid concentration and variability (RSD) (Fig. 5G). Interestingly, we observed decreasing metabolic variability for stronger phenotypic alterations (wt > *leu4Δ* > *bat1Δ* > *oac1Δ*). This suggests that genetic perturbations promoted the loss of metabolic flexibility. In addition, it was observed that *oac1Δ* amino acid concentration was lower in general. With the exception of phenylalanine and tyrosine, which originate from chorismate ^22^, all other amino acids had lower concentrations upon *oac1Δ* deletion that in the wild type.

We further tested whether the amino acid concentration variability was dependent on the growth phase, but it was in fact stable across growth phases (Fig. 5H). However, this variability was generally higher in the wild type and wild type-like strains, *bat2Δ* and *leu9Δ,* and lower in the remaining strains, that affected the most the protein profile of the BCAA pathway and the general metabolic landscape. This indicates that, on the one hand, although the diauxic shift is a transition phase it does not rely on increased metabolic concentration variability. On the other hand, we can speculate that the reduced concentration variability observed upon genetic perturbations could jeopardize the overall variability of the population and thus, the overall population adaptability to stresses caused by a wide range of glucose concentrations or different environmental conditions.

### The leucine branch controls respiration enzymatically, while the shared branch relies on other mechanisms

Next, we asked to what extent the metabolic fluxes are remodeled during diauxic shift. As direct experimental quantification of intracellular fluxes is not feasible, we utilized the protein profile of the BCAA pathway, along with metabolomics data, to constrain a thermodynamically consistent model developed by Oftadeh O. *et al* in 2021. The *S. cerevisiae* yETFL.cb model was based on the Yeast8 genome-scale, and was constrained to eliminate thermodynamically infeasible fluxes, while the feasible ranges for metabolite concentrations and reaction thermodynamic displacement were defined ^23^ . Our genetic background was imposed in the form of gene deletions. The growth medium was adjusted and supplemented with the respective auxotrophic metabolites. We extended the model with additional reactions describing the mitochondrial transport of BCAA and their respective keto acids by adding degrees of freedom to the model.

To further constrain the model, we integrated experimental data, specifically the protein levels of the BCAA pathway and intracellular metabolite concentrations categorized across various metabolic pathways. The protein abundances were integrated into the model in the form of ratios relative to Leu1 protein, which was the most abundant enzyme in both the yETFL.cb model and in the experimental data (Fig. 3A, Model). The mean absolute concentrations of the 13 metabolites were directly integrated into the model, and the relative metabolite concentrations were included in the form of metabolite concentration ratios considering the maximal metabolite concentration of the compartment where the reaction was assigned (Materials and methods). In all cases the estimated standard errors were integrated in the form of bounds in the metabolite values. For the rest of the proteins and metabolites we used the parameters defined in the yETFL.cb model. We refer to the data-unconstrained and data-constrained models as REF and WT models, respectively.

For the characterization of the fluxomic space, we employed thermodynamics-based flux variability analysis (TFVA). This method explores the feasible flux ranges, identifying their minimum and maximum values. We evaluated approximately 350 reactions involved in amino acid metabolism, central carbon metabolism (Glycolysis, TCA-cycle, glyoxylate cycle), and transport reactions at the characteristic time points that hold the steady state assumption, namely T2 and T6. The experimental data integration reduced the thermodynamic space of the network, narrowing the range of possible physiological states the cell can adopt. Simulations for growth optimization revealed that although both models fermented at their maximum growth rate (REF 0.44h-1 and WT of 0.37 h-1) only the WT model fermented at 0.37 h-1 (Fig. S10). At T2, the glucose consumption and ethanol secretion fluxes assumed higher rates in the WT than in the REF model, whereas at T6 similarly low levels of these fluxes were observed in both models. This indicates that both models operate in a respiratory regime at T6 (Fig. 6A upper left). Hence, the sole integration of the WT T2 experimental data, without ad hoc modifications in the model, lowered the maximum growth rate and the Crabtree effect onset. Such behavior resulted from the fact that the glycolytic flux ranges attained higher rates in the WT than in the REF model. In agreement with Zampar *et al.*^24^ we observed that while the high glycolytic fluxes decreased at T6, TCA cycle fluxes were similar at both physiological states (Fig. 6A). This resembles the second temporal step towards the gluconeogenic regime in yeast - the onset of the glyoxylate cycle - when most of the remaining glucose in the growth medium is respired, just before ethanol flux reversal is established and its consumption starts.

**Figure 6.**
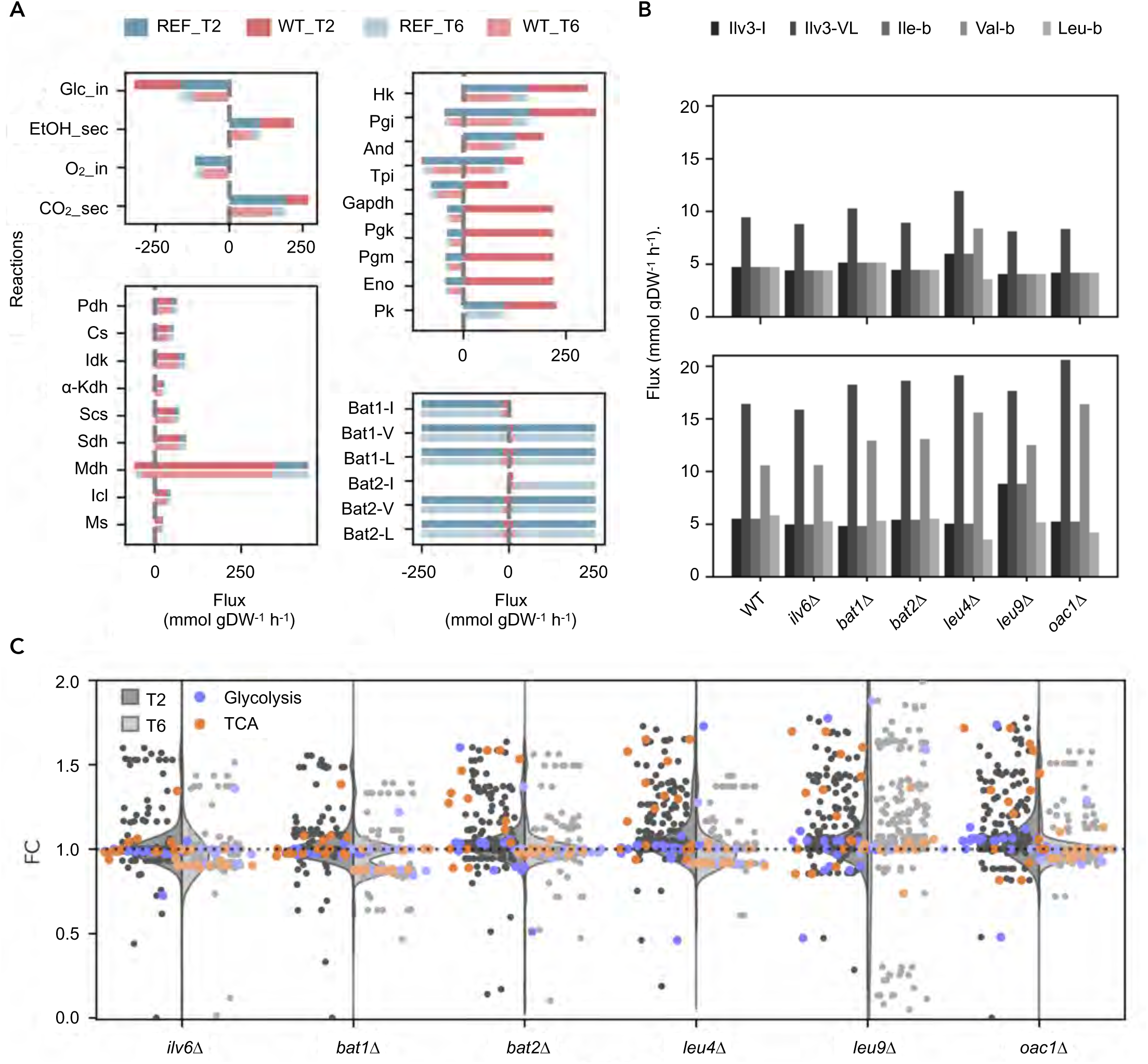
Comparative analysis of thermodynamic flux variability in BCAA and in central carbon metabolism across growth phases. **A)** Thermodynamic flux variability analysis. **B)** Flux distribution in BCAA pathway at the T2 uppper and T6 botom. Names of the pathway branches correspond to the following segments of the pathway Ile_shared-from Thr to KMV; Val&Leu_shared - from PYR to KIV; Ile_b from KMV to Ile; Val_b from KIV to Val and Leu_b from KIV to Leu. Values represent the average of the maximal and minimal value of the flux through each unidireccional reaction and, and the shared fluxes Val_b and Leu_b were split based on their proportions (Material and methods). **C)** Overview of metabolic flux distribution across central carbon and amino acid pathways. Reactions are colored by pathway. Values represent the fold change of average of the maximal and minimal value of the flux through each reaction in each deletion strain relative to the WT.

A more detailed examination of the WT model reveled that the BCAA fluxes were limited to about half the range allowed in the REF model, at both T2 and T6 (T2= 0.46+/-0.16 and T6=0.63+/-0.14) with the exception of Bat1 and Bat2 transaminases and their respective keto acid transporters, for which the flux range was 10 times lower in the WT than in the REF model. In both models the BCAA pathway reactions were mostly unidirectional except for the BCAA transaminases and BCAA transport reactions that were mostly bidirectional. This pattern revealed that the BCAA pathway can display different flux topologies i.e. valine and leucine could be synthesized either in the mitochondria or the cytoplasm (Fig. 6A bottom left, Supp Data 4). Additionally, due to the capacity of these transaminases for generating internal cycles, loops where metabolites are cycled without a net input or output — meaning the fluxes through those reactions can become unrealistically high without violating mass balance. Thus, the BCAA transaminases and BCKA transport reactions displayed the highest flux rates of the pathway, which did not match the overall pathway flux. Therefore, these redundant reactions were excluded when estimating the pathway’s flux remodeling.

The overall pathway flux distribution was estimated by the average of the maximal and minimal flux values through each of the unidirectional reactions. The shared branch of the pathway uses the same enzymes, but depending on the substrates it can produce KMV that leads to Ile, or KIV that is used to produce Val or Leu. The fluxes producing isoleucine were directly determined based on their average flux. Valine and leucine fluxes were considered complementary and collectively accounted for all the flux passing through the shared enzymes of the pathway that used the intermediates specifically for KIV production. We observed that the fluxes through the pathway’s shared enzymes were similar at T2 for the three BCAA. However, at T6, fluxes leading to KIV displayed a twofold flux increase, that was mostly funneled towards valine rather than to the leucine committed branch (Fig. 6B). This observation implies that only the reactions involved in valine production undergo adaptation to meet metabolic demands during this fermento-respiratory transition.

We then examined how genetic perturbations affected the BCAA pathway at the flux level. At T2, the *leu4*Δ mutant exhibited the most distinct flux distribution compared to WT, with a 25% increase in flux through the shared branch of the pathway, primarily at the expense of a similar reduction in flux through the leucine branch. By T6, all knockouts of the leucine-committed branch (*leu4*Δ, *leu9*Δ, and *oac1*Δ) displayed similar changes, higher fluxes towards valine and lowers through the leucine committed branch. However, while isoleucine fluxes displayed WT levels in *leu4*Δ and *oac1*Δ strains, they were elevated in the *leu9*Δ knockout (Fig. 6B). The flux patterns observed in *leu4*Δ and *oac1*Δ strains were consistent with their metabolic profile (Fig. 5A), suggesting that either upregulation of the Leu9 protein in these mutants (Fig. 3E) is insufficient to support WT fluxes through the leucine-committed branch, or that Leu1 downregulation restricts flux through this branch. Finally, the remaining knockouts did not significantly alter BCAA pathway fluxes. This suggests that in the *bat1*Δ mutant, the flux range assumed by the WT BCAA pathway can accommodate the metabolite concentration changes observed in this strain without substantially shifting metabolic fluxes (Fig. 5A and 6B).

We continued to broaden our scope towards the remaining tested fluxes, which represented other amino acid and central carbon metabolic pathways (Fig. 6C). As in the case of the BCAA pathway, we observed more striking changes in the general fluxomic space at T6 than at T2 (Fig. 6B-C). In particular, the flattened distribution of *leu9*Δ knockout stands out. Sucha distribution was due to a flux reduction in transport reactions between compartments, mainly amino acid transport between the cytoplasm and vacuole, and to an increased glucose intake, that boosted fluxes through central carbon metabolism. This strong remodeling was not detected by our longitudinal statistical test (Fig. 5B and Supp Data 3), but it was detected by the PCA analysis. *leu9*Δ was close to the WT at early time points, but as the diauxic shift approached, it moved away, eventually becoming the most distant KO from WT at T6 as detected by PC1 (Fig. 5F and Fig. S9). This finding highlights the increasing relevance of generating multi-omics approaches with spatial resolution at the subcellular level.

During logarithmic growth, we found that the fold change in fluxes for each strain compared to WT was centered around 0, indicating that, at this early point, the metabolic fluxes in the mutant strains remain largely like those in the WT. However, the knockouts of the leucine-committed branch (*leu4*Δ, *leu9*Δ, and *oac1*Δ) and bat2Δ strain, that is also involved in leucine biosynthesis, displayed an increase of up to 75% in TCA-cycle fluxes as well as in the isocitrate lyase Icl1, a key reaction of the glyoxylate cycle. The flux increases through these typically glucose repressed pathways, was the product of rerouting AcCoA from the BCAA Leu4 Leu9 reactions to TCA-cycle via Pyruvate dehydrogenase (PDH), in a glycolytic independent manner (Fig. 6C).

In contrast, at T6 *ilv6,* bat1Δ, *leu4*Δ, and *oac1*Δ displayed fold change distributions skewed to values below 0. This general decrease of the fluxes relative to the WT included a 15%-10% flux drop on glycolysis, TCA cycle and Icl1 isocitrate lyase as well as reactions involved in other amino acid pathways. Such a general response could not be traced back metabolically to the BCAA pathway, because it affected intake reactions. The observed behavior suggested that at this later timepoint metabolic adjustments in response to the mutations have underlying mechanisms that relay in other biological layers, such as transcriptional and posttranslational. These modifications usually take time to manifest, resulting herein modest, but broad differences (Fig. 6C).

## DISCUSSION

The comprehensive exploration of the leucine metabolic pathway has gained importance due to increasing evidence purposing this pathway as a fundamental link between amino acid and central carbon metabolism during yeast diauxic shift^10^. As a building block for protein synthesis, leucine assumes a critical role in amino acid metabolism and biomass production. The control that leucine and Bat1 exert on the TORC1 pathway^10^ extends the role of the BCAA pathway to the transition from logarithmic to stationary growth^8^. Recently Interestingly, TORC1-leucine and valine mediated activation has recently also been proposed as a metabolic cell cycle control mechanism^25^. Despite substantial evidence placing the BCAA pathway as a primary link between energy and biomass production, it is neither clear how leucine or the BCAA pathway influence the diauxic shift, nor how the BCAA pathway itself is regulated. Here, we present a comprehensive study on the dynamic behavior of the BCAA pathway that combines experimental and computational methods to explore the behavior of the metabolic landscape throughout the diauxic shift in the wild type and upon the systematic deletion of the regulatory elements of the pathway.

We demonstrated that in yeast the BCAA pathway behaves in two separate branches with opposing behavior throughout diauxic shift. The leucine branch exhibited preferential activity during fermentative growth, characterized by energy production through glycolysis and high growth rates^5^. In contrast, the shared-branch displayed higher activity when the respiratory regime is established, in which energy production relies increasingly on respiration and reduced growth rates are observed^5^ (Fig. 3A-C, 4B and G). The divergence between these two profiles aligns with reported TORC1 activity levels^8–9^, and was not related to the sub-cellular localization of the enzymes involved in each branch (Fig. 1A and 2A). Implying an association with their specific metabolic functions rather than a fully compensatory response to mitochondrial volume increase. In fact, while the shared-branch protein profile counteracted the dilution of this branch, the mitochondrial volume increase magnified the drop in Leu-branch protein concentration while growth phases progresses^11, 15, 12^ ^and^ ^26^ (Fig. 3A, 2B-D and G).

We further explored the regulation of the BCAA pathway during the yeast diauxic shift by examining the pathway response across all prototrophic BCAA pathway deletion strains. At the protein level, we found that the elements responding to genetic perturbations—Ilv2, Bat2, and Leu9—differed from those regulating the pathway’s response, namely *ilv6Δ*, *bat1Δ*, *leu4Δ* and *oac1Δ*. Although responsive and regulatory elements participated in the same reactions (Ilv2/Ilv6, Bat2/Bat1 and Leu9/Leu4), the responsive elements not only reacted to the absence of its pair, suggesting a pathway-specific rather than a partner-specific response^27^. In particular, the response of Bat2 and Leu9 upon the deletion of their paralogous partners revealed that, while these responsive elements could substitute catalytically the absence of their paralogous pair, they could not substitute their regulatory role.

The systematic deletion of the regulatory elements of the pathway allowed us to individually modify the strength of each feedback loop of the pathway and, thus, establish which regulatory loop(s) governed the pathway under the biosynthetic conditions tested. Considering that i) *ilv6Δ* knockout rendered the pathway unresponsive to valine inhibition due to the loss of the Ilv2/Ilv6 regulatory subunit ^28^; and ii) that *bat1Δ* deletion decreased the strength of the valine negative feedback loop by lowering valine intracellular concentration^29^ (Fig. 5A), one would expect that completely removing the loop (*ilv6*Δ) has a greater effect than merely lowering the inhibitor (*bat1*Δ). However, the greater metabolic response to *bat1Δ* deletion compared to *ilv6Δ* strain (Fig. 3E, 5A-C and 6C) indicates on the that the Val inhibition feedback loop has a minor role under the biosynthetic conditions tested, and that on the other hand Bat1 has an additional role beyond feedback inhibition.

Analysis of the protein and metabolic response upon *bat1*Δ, revealed that the extended metabolic effect upon *bat1Δ* deletion associated this strain with a more intricate phenotype (Fig. 5B-F). The simultaneous reduction of valine and isoleucine alongside the accumulation of cis-aconitate and α-ketoglutarate during log phase, suggested that Bat1 does not couple the BCAA pathway to the TCA-cycle through the BCAT-mediated transamination of α-ketoglutarate, but instead it directly interacts with the TCA-pool. In agreement, the TFVA suggests that the accumulation of cis-aconitate and α-ketoglutarate is alleviated, when respiratory metabolism is established. This occurs by a decrease in the fluxes through energy producing pathways (glycolysis, TCA- and glyoxylate-cycle), allowing the BCAA to drain metabolites towards leucine biosynthesis. This mechanism is consistent with the synthesis of leucine from glucose breakdown via pyruvate and acetyl-CoA^25^, as well as with the Bat1- Aco1 complex disruption phenotype that blocked the TCA cycle at the step of pyruvate and acetyl-CoA incorporation^10^. Taken together, we propose that Bat1 controls TCA cycle flux through its physical interaction with Aco1, an effect that becomes more evident in later growth phases when cells increasingly rely on respiratory pathways for energy and biomass production. This results in a response that extends to glycolysis, potentially by transcriptionally reprogramming the respiratory transition.

We also perturbed the leucine negative feedback loop. Previous studies have indicated that leucine feedback loop is weakened by the deletion of *leu4Δ*^13^. The lack of Leu4 facilitates the assembly of the Leu9-Leu9 leucine-resistant isoform, leading to leucine and α-IPM accumulation and the subsequent Leu3-α -IPM dependent activation of *LEU9.* We observed that despite Leu9 was up-regulated upon *leu4Δ* deletion (Fig. 3E), leucine levels were consistently lower than in the wild type (Fig. 5A). Unlike in previous studies, we observed a down-regulation of the downstream element Leu1^30^. Although statistical methods relying on mean or variance might have missed its significant effects due to its bimodal distribution and associated high variability (Fig. 3E), Leu1 displayed a strong response, which in turn explained the decrease in leucine levels and the up-regulation of Leu9 (Figs 3E and 5E). Similar to the results for *bat1*Δ, low leucine levels were accompanied by an increase in cis-aconitate and α-ketoglutarate. However, unlike in the *bat1*Δ strain*, in-silico* evidence supported that the blockade of the leucine committed branch (*leu4Δ, leu9Δ*, *oac1Δ* and *bat2Δ*) re-routed pyruvate from the leucine committed branch towards TCA- and glyoxylate-cycle, via PDH, resulting in leucine depletion and accumulation of cis-aconitate and α-ketoglutarate. Such flux reconfiguration highlights that BCAA leucine committed branch governs respiratory metabolism through its enzymatic activity ^31, 32^ during the fermentative phase.

When respiratory metabolism was established, we observed that, similar to the *bat1*Δ strain, the *leu4*Δ knockout showed decreased fluxes through glycolysis, TCA and glyoxylate-cycle. In neither of these strains the effect could be directly traced back metabolically, suggesting the involvement of additional regulatory levels, such as transcriptional reprogramming. In agreement, despite their opposite effects on leucine levels, leu4Δ led to decreased leucine, while *bat1*Δ, due to the absence of *Leu1* downregulation, resulted in increased leucine. Fluxes towards α-IPM were elevated in both cases and, in consequence, both strains exhibited *LEU9* up-regulation. Additionally, *Bat2* upregulation in both strains align with its previously described induction via Gcn4 under amino acid deprivation, which in this case is triggered by low Ile and Val levels in *bat1*Δ and low Leu levels in *leu4*Δ. Therefore, the response observed in these knockout strains at later growth phases is likely the result of at least two transcriptional programs i.e. α-IPM-Leu3 activation due to high α-IPM levels and Gcn4 activation, driven by low levels of Ile and Val in *bat1*Δ, and Leu in *leu4*Δ.

In contrast, the *oac1Δ* strain does not show a similar behavior to the one observed in *bat1Δ* and *leu4Δ* at T6. Similar to the previously described *leu3*Δ induced amino acid deprivation, in the *oac1Δ* strain we observed low amino acid levels in general, and lack of Leu9 up-regulation, indicating that *oac1Δ* deletion leads to low levels of α-IPM and thus poor Leu3 activation. This amino acid deprivation is consistent with the observed Bat2 up-regulation, that as mentioned above *BAT2*-induced expression through Gcn4 is a response to amino acid deprivation. Considering that TFVA indirectly indicates that the perturbation of the KIC negative feedback alone does not fully explain the extended impact of *oac1Δ* over metabolism, we propose that the response observed in the *oac1Δ* knockout is the result of the lack of activation due to low α-IPM-Leu3 levels, that drive general low amino acid levels resulting in Gcn4 activation^19^ ^and^ ^33^ .

Taking together, our experimental approach revealed that the valine feedback loop has a minor effect compared to leucine inhibition. Initially, leucine regulation operates through direct metabolic effects; however, at later time points, its impact—like other pathway perturbations—shifts from metabolic control to compensatory transcriptional programs. This is evident in strains such as *bat1*Δ and *oac1*Δ, where the response is not directly driven by metabolic disruption but likely by the activation of regulatory networks. Although we demonstrated here the relevance of the temporal resolution, there is still work to do not only to integrate other biological layers such as the transcriptional response, but also the subcellular spatial resolution.

Finally, the wildtype temporal pattern of the pathway and the generally coordinated metabolomic landscape support the assumptions that global physiological factors such as growth rate and nutrient availability govern metabolism through global gene programs. Furthermore, the overlap of the affected metabolic pathways upon genetic perturbations supports the finding that the BCAA pathway is a functional-metabolic cluster ^34^. In particular, the extended impact and wide metabolic overlap between *bat1Δ, leu4Δ* and *oac1Δ* suggest the action of a GCN4 global program, while *bat1Δ* and *leu4Δ* potentially rescued that effect by accumulating α-IPM. However, specific metabolic signatures, in which the metabolites affected within each pathway did not overlap, challenge the modular assembly of metabolic pathways in structural units with related functions and the idea that few hub metabolites dominate metabolism ^35^ ^and^ ^36^

## MATERIAL AND METHODS

### EXPERIMENTAL MODEL AND DETAILS

#### Strains and Media

BCAA pathway deletion strains were constructed in the S288c derivate Y8205 that constitutively express mCherry (strain background*: Matα Pdc1pr_mCherry-Ca3URA ho3*Δ*0 can1::ste2(pra)_Sp_his5 lyp1::*direct repeat)^37^. The open reading frame of each gene of the pathway was substituted by the *NATMX4* resistance cassette (*xxxΔ::NAT*) by PCR-based homologous recombination. Since the query collection included the neutral-insertion *ho*Δ, a *hoΔ*::*NATMX4* strain was generated, this unrelated deletion background was used as wild-type control. The *NATMX4* resistance cassette was amplified from pAG25, primer sequences for gene-knockout and confirmation PCR were obtained from the Yeast Deletion Project^38^. Strains and primers are shown in Supp_Table_S1.

To generate the GFP protein fusions the HIS prototrophy marker of the GFP fusion collection (derived from the ATCC 201388, MATa his3Δ1 leu2Δ0 met15Δ0 ura3Δ0) Invitrogen was substituted by direct gene replacement with the *kanMX4* cassette (*XXX-GFP-KAN*)^39^. The *kanMX4* resistance marker was obtained from the pFA6a plasmid by Not1 enzymatic digestion. The *leu2Δ* auxotrophy selection marker was substituted with the *LEU2* wt gene by PCR-based gene replacement, *LEU2* wt gene was amplified form ho*Δ::NAT strain* (Y8205 background) with the primers P25 and P26 (Supp_Table_S1). The protein fusion strain *BAT1-GFP-KAN* was constructed *de novo* by PCR-based homologous recombination. The *GFP-KAN* module was amplified form the *ILV2-GFP-KAN* strain (*ATCC 201388* background) with the primers P27 and P28 (Supp_Table_S1). All strains were confirmed by PCR using primers A and kanB from the Yeast Deletion Project Supp_Table_S1.

The following growth media were used: i) Yeast-extract peptone with 2% dextrose (YPD). ii) Synthetic-complete medium (SC) was 6.7 g/L yeast nitrogen base without amino acids and 2% glucose, amino acids to satisfy auxotrophic requirements (methionine, histidine, uracil and methionine) were added at 0.01% w/v (-Cold Spring Harbor Laboratory Manual 2005). Antibiotic-resistant strains were grown in media supplemented with 100 μg/ml nourseothricin (ClonNAT, Werner BioAgents), 200 μg/ml G418 (Invitrogen). Ammonium was replaced by 1g/L monosodium glutamic acid in SC medium with antibiotics.

#### Generation of yeast libraries

Collections of GFP-tagged proteins in wild-type and knockout backgrounds were generated by synthetic genetic array methodology (SGA). The GFP-tagged strains were mated to an array of 144 Y8205-mCherry BCAA pathway single-deletion starter strains. This mating step was followed by diploid selection, sporulation, and three rounds of haploid selection (*HIS^+^* for *MATa* mating type, *G418^+^*for fluorescence marker, and *clonNAT* for knockout selection). The resulting phototrophic strains were (*MATa PDC1-RFP-CaURA3MX4 can1*Δ*::STE2pr-SpHIS5 lyp1*Δ *ura3*Δ*0 his3*Δ*1 LEU2 MET15 xxx*Δ*::natMX4 XXX-GFP-kanMX4).* Likewise, the CFP-labeled reference strain was obtained from the YEG01-CFP starter strain were also generated by the same procedure of combining ho neutral deletion with all GFP backgrounds. The libraries were constructed in quadruplicate and Colony and transferred manually with a 384-head pin tool (V&P Scientific, VP384F); antibiotic concentrations used for selection were 200 µg/ml G418 (Invitrogen) and 100 µg/ml clonNAT (Werner BioAgents). In the following only phototrophic strains were used.

### HIGH-THROUGHPUT FLOW CYTOMETRY

#### Preparation of co-cultures

Each library was grown individually to saturation in YPD medium 96-well plate format. Medium was dispensed (600 µl) with a MicroFill Microplate Dispenser (BioTek) onto 1.0-ml polypropylene plates (Nunc 260251), and cultures were incubated in a Multitron Infors platform shaker at 30°C with shaking at 999 rpm. Each experimental run involves the co-culturing of wild type and knockout libraries-wt (GFP-tagged protein in CFP background) and ko (GFP-tagged protein in a deletion background constitutively expressing mCherry).

The two saturated libraries were mixed in a 1∶1 ratio to a final volume of 150 µl in 96-well plates (Corning 3585). Fresh SC media (600 μl) in deep-well plates (Nunc 260251) with disposable plastic covers was inoculated with 60 μl of saturated co-cultures using an automated robotic station (Tecan Freedom EVO200) that integrates a plate carrousel (Liconic STX110), a high-performance multilabel plate reader (Tecan Infinite M1000), a 96-channel pipetting head, an orbital plate shaker, and a robotic manipulator arm, all contained in an isolated environmental. Co-cultures were maintained at constant temperature (30°C) and relative humidity (70%) with constant shaking; Strains were then grown to early-log phase (∼10 h) and followed every two hours up to stationary-phase (10 h) . To analyze the libraries, cells were first transferred into 96-well plates with TE (10 mM Tris and 1 mM EDTA pH 8) by automated pipetting. After transferring the plates to a plate reader to collect data for optical density at 600 m (OD_600nm_), plates were analyzed in a flow cytometer.

#### Flow Cytometry: Instrumentation, acquisition and data preparation

A flow cytometer with a high-throughput autosampler (BD LSR Foressa X20) was used to record fluorescence from GFP, CFP, and mCherry fluorophores. Optimal voltage was adjusted using single stained control strains expressing only one fluorophore. GFP was excited with a 488-nm laser, and fluorescence was collected through a 530/30 band-pass and 550LP emission filter. CFP was excited with a 405-nm laser, and fluorescence was collected through a 525/50 band-pass filter and a 505LP emission filter. mCherry was excited with a 561-nm laser, and fluorescence was collected through a 610/20 band-pass and a 640LP emission filter. Cells were measured in high-throughput mode at a flow rate of 0.5 µl/s for 8 s.

A Matlab workflow was developed to enable an automated analysis of data, fcsread.m (Robert Hanson, available at Matlab central) and scatter_kde (Haëntjens, 2018) functions were used to import fcs raw and to scatter plot with a probability density estimate. For each well (co-culture), analysis was as follows steps: 1) Remove cell doublets by manually defining a gate based on the forward scatter of the populations (FSC_H and FSC_W). 2) Remove cell debris using a gate based on the forward and side scatter of the populations (FSC_A and SSC_A). 3) Compensate GFP, CFP, mCh channels ^40^ using single stained color controls to estimate the spillover. The mean of the spillover was considered the spillover constants of the compensation matrices. 4) Classify the cells into mCh expressing or CFP expressing cells defining a gate-based CFP and mCh recorded levels in single stained controls. 5) Volume normalization was obtain dividing the GFP fluorescent signal by FSC_A.

#### Fluorescent microscopy

To confirm subcellular localization of GFP tagged strains, cells were stained with MitoTracker Red CMXRos (Molecular Probes) according to manufacturer’s specifications. Co-localization between the MitoTracker and GFP was determined through sequential imaging. Confocal images were obtained using a FluoView FV1000 laser confocal system (Olympus) attached/interfaced to an Olympus IX81 inverted light microscope with a 60x oil-immersion objective (UPLASAPO 60x O NA:1.35), zoom x20.0 and 3.5 μm of confocal aperture. The excitation and emission settings were as follows: GFP excitation at 488 nm; emission 520 nm BF 500 nm range 30 nm; MitoTracker excitation 543 nm; emission 598 nm, BF 555 nm range 100 nm. The subsequent image processing was carried out with Olympus Fluo View FV1000 (version 1.7) software.

### UNTARGETED METABOLOMICS

#### Chemicals

LC-MS grade water, acetonitrile and methanol were obtained from Th. Geyer (Germany). High-purity ammonium acetate, ammonium formate, acetic acid and formic acid were purchased from Merck (Germany). Stable isotope labelled amino acids (MSK-A2-1.2; Cambridge Isotope Laboratories, MA, USA) were used as internal standards for untargeted metabolomics.

#### Metabolite extraction

Cells were collected by rapid centrifugation for 1 min at 5000 rpm at 4°C with a 5415R microcentrifuge (Eppendorf, Hamburg, Germany), washed with PBS and re-centrifuged, the medium discarded and the samples were frozen in liquid nitrogen and stored at −80°C for later metabolite extraction. Metabolite extraction was performed by addition of acetonitrile:methanol:water (2:2:1, v/v) to the yeast pellet samples. The added volume was adjusted to a uniform OD_600nm_ of 4 and 5 for batch 1 and 2, respectively. Subsequently, samples were vortexed thoroughly for 10 min, ultrasonicated on ice for 5 min, and incubated at -80°C for 20 min. After centrifugation for 10 min at 15,000 × *g* and 4 °C with a 5415R microcentrifuge (Eppendorf, Hamburg, Germany), samples were spiked with internal standard mixture in acetonitrile:methanol:water (2:2:1, v/v). The final samples were transferred to analytical glass vials and the LC-MS/MS analysis was initiated within two hours after the completion of the sample preparation.

#### LC-MS/MS analysis

LC-MS/MS analysis was performed on a Vanquish UHPLC system coupled to an Orbitrap Exploris 240 high-resolution mass spectrometer (Thermo Fisher Scientific, MA, USA) in positive and negative ESI (electrospray ionization) mode. Chromatographic separation was carried out on an Atlantis Premier BEH Z-HILIC column (Waters, MA, USA; 2.1 mm x 100 mm, 1.7 µm) at a flow rate of 0.25 mL/min. The mobile phase consisted of water:acetonitrile (9:1, v/v; mobile phase phase A) and acetonitrile:water (9:1, v/v; mobile phase B), which were modified with a total buffer concentration of 10 mM ammonium acetate (negative mode) and 10 mM ammonium formate (positive mode), respectively. The aqueous portion of each mobile phase was pH-adjusted (negative mode: pH 9.0 via addition of ammonium hydroxide; positive mode: pH 3.0 via addition of formic acid). The following gradient (20 min total run time including re-equilibration) was applied (time [min]/%B): 0/95, 2/95, 14.5/60, 16/60, 16.5/95, 20/95. Column temperature was maintained at 40°C, the autosampler was set to 4°C and sample injection volume was 6 and 5 µL for batch 1 and 2, respectively. Analytes were recorded via a full scan with a mass resolving power of 120,000 over a mass range from 60 – 900 *m/z* (scan time: 100 ms, RF lens: 70%). To obtain MS/MS fragment spectra, data-dependent acquisition was carried out (resolving power: 15,000; scan time: 22 ms; stepped collision energies [%]: 30/50/70; cycle time: 900 ms). Ion source parameters were set to the following values: spray voltage: 4100 V (positive mode) / -3500 V (negative mode), sheath gas: 30 psi, auxiliary gas: 5 psi, sweep gas: 0 psi, ion transfer tube temperature: 350°C, vaporizer temperature: 300°C.

All experimental samples were measured in a randomized manner. Pooled quality control (QC) samples were prepared by mixing equal aliquots from each processed sample. Multiple QCs were injected at the beginning of the analysis to equilibrate the analytical system. A QC sample was analyzed after every 5^th^ experimental sample to monitor instrument performance throughout the sequence. For determination of background signals and subsequent background subtraction, an additional processed blank sample was recorded. Data was processed using MS-DIAL^41^ and FreeStyle 1.8 SP2 (Thermo Fisher Scientific) and raw peak intensity data was normalized via total ion count for relative metabolite quantification. Feature identification was based on accurate mass, isotope pattern, MS/MS fragment scoring and retention time matching to an inhouse library (EMBL-MCF 2.0, please cite, DOI: 10.1007/s11306-024-02176-1).

### QUALITY CONTROL AND DATA CORRECTION

#### Growth phase correction and batch normalization

In order to compare the phenotype of different strains, displaying different growth rates, at the same growth phase, mean growth phase obtained from biological triplicates were shifted to align the growth phases of all strains. Consensus time points spanning early logarithmic phase to post diauxic shift were established and metabolomics data was shifted accordingly. Key central carbon metabolites (known to respond to the diauxic shift) were used to confirm qualitatively the proper growth phase alignment.

Samples were processed in two different batches; each one contained a technical triplicate of the wild type strain. Wild type from batch one was referred as anchor sample and wild type in batch two served as reference strain. Target metabolites from batch two data set were normalized to be on the same scale as their counterparts in batch one by computing a scaling factor, the ratio of the mean intensity of the anchor samples to the reference set, then the computed scaling factor was used to multiply the values on the target set (batch 2 set).

#### Metabolomics data preparation

Corrected data was subjected to different transformations depending on the purpose of the analysis. For cross correlation and enrichment analysis, the data was log transformed, for visualization purposes the data was scaled with respect to the first time point. For statistical MB-test data was log transformed and autoscaled^42^.

#### Amino acid intercellular estimation

Intracellular concentration of free amino acids in wt, *leu4Δ* and *leu9Δ*; was determined using the median OD_600nm_ of the analyzed samples, the approximate extraction volume of 110 µl and considered that 1 OD = 3.2*10^7 cell/ml and the cell volume as 45.45 fL. Since each batch contained the wt strain, we used amino acid concentration in these strains and the relative changes in the knockout strains to estimate the free amino acids in the remaining strains. As controls we used *leu4Δ* and *leu9Δ* intracellular concentrations and the estimated intracellular concentrations.

### DATA ANALYSIS

#### Multivariate empirical bayes statistic (MB-test)

We implemented two-sample multivariate empirical Bayes statistics (the MB-statistics)^43^ to rank the response of protein or metabolites to different deletions in order of interest from longitudinal replicated high-throughput flow cytometry or LC/MS untargeted metabolomics time course experiments. The statistical test was performed using the R package “time course” ^44^ . A p-value for the change in the overall protein and metabolic profile was calculated from the MB-statistic using the F-statistic cutoff values were p < 0.01 for proteomics and p < 0.05 for metabolomics.

#### Principal Component Analysis

Protein abundances measured during diauxic shift in the wild type and lockout strains were auto-scaled and used as independent variables in the loading plots for construction of the principal component analysis. K-mean clustering was set to partition the 35 observations into the maximum number of clusters that are not overlapped.

#### Cross-correlation analysis

Cross-correlation was applied to metabolite intensity and the cluster of metabolites were defined by sharing mean average cross-correlation values above 0.75 with all the components of the cluster.

#### Metabolite enrichment and pathway impact analysis

Metabolite enrichment analysis (MEA) was performed with MetaboAnalystR Package (https://www.metaboanalyst.ca/docs/About.xhtml). Metabolites identifiers against S.*cerevisiae* KEGG pathways were used to assess pathway overrepresentation. Statistical significance was based on the Holm statistic associated adjusted p-value < 0.05. Edges between metabolic pathways were displayed when two pathways shared more than 10% of their elements, the weight of the edges reflected the fraction of metabolites shared with respect to the larger metabolic pathway. Impact and Importance analysis were conducted in MetaboAnalyst 3.0 (Xia et al. 2009, 2015). Pathway impact Statistical significance was based on the Holm statistic associated adjusted p-value < 0.05.

#### Update and constrain the genome scale model

We utilized the yETFL.cb model, an extended genome-scale model based on Yeast8, which integrates expression and thermodynamics-based constraints. The model was adjusted to reflect our experimental conditions, mitochondrial transport of branched-chain amino acids (BCAAs) and their respective keto acids, as well as additional gene complexes for threonine deaminase and 2-isopropylmalate synthase were incorporated into the model. Knockout models were generated by implementing the respective gene deletions including the *met15* knockout background we worked with.

The absolute concentrations of 13 metabolites was integrated to the model directly and the relative concentrations of 64 metabolites were integrated as relative to the compartment define maximum abundance concentration considering the maximum abundance over the experimental time course. Protein abundance ratios were normalized to Leu1 experimental measurements and these relative abundances were integrated to the model as considering the Leu1 original yETFL.cb model concentration. Standard errors were applied as bounds for metabolite and protein concentrations, and remaining parameters were retained from the original yETFL.cb model.

#### Model simulation and thermodynamics flux variability analysis

We performed thermodynamics-based flux variability analysis (TFVA) using the pyTFA^45^, a python package implementation for Thermodynamics-based Flux Analysis that aids the determination of feasible flux ranges under steady-state conditions. Flux simulations were conducted at two key time points (T2 and T6), evaluating approximately 350 reactions spanning amino acid metabolism, central carbon metabolism, and transport processes. Model optimization was carried out for maximal growth rate to assess metabolic shifts between fermentative and respiratory states.

## AUTHORS CONTRIBUTIONS

Project conceptualization: XE-F. Supervised and planned experiments/data analysis XE-F and EK. Library strain construction XE-F. Physiological measurements XE-F. Protein profile XE-F, OB. Metabolomics XE-F, OB and BD. Model curation and data integration XE-F and OB. Original writing XE-F. Supported the work and helped with the writing AL, TA and EK. Contribution with reagents and funding AL, TA and EK. Funding acquisition: XE-F. Project supervision: XEF, TA and EK. All authors have read and approved the manuscript.

## AKNOWLEDGMENTS

We thank Dr. Diana Ascencio for her support on high-throughput flow cytometry measurements. X.E.-F. was supported with a postdoctoral grant from CONACYT (CVU 420248)

## DISCLOSURE AND COMPETING INTEREST STATEMENT

The authors declare no competing interests.

## FIGURE LEGENDS

**Figure Synopsis:** Integration of BCAA pathway protein profile and metabolomic context into a thermodynamically curated genome-scale model identifies pathway dynamics and regulation at different growth phases and highlights role of the BCAA pathway in metabolic network integration during the diauxic shift.

`o` The BCAA pathway operates in two branches: the leucine-committed branch shows a fermentative profile aligned with TORC1 activity, while the shared segments display a fermento-respiratory signature.
`o` Different feedback loops control specific sections of the pathway and operate at different growth phases.
`o` Misregulation of the pathway result in altered flux distributions in distant metabolic pathways including central carbon metabolism.

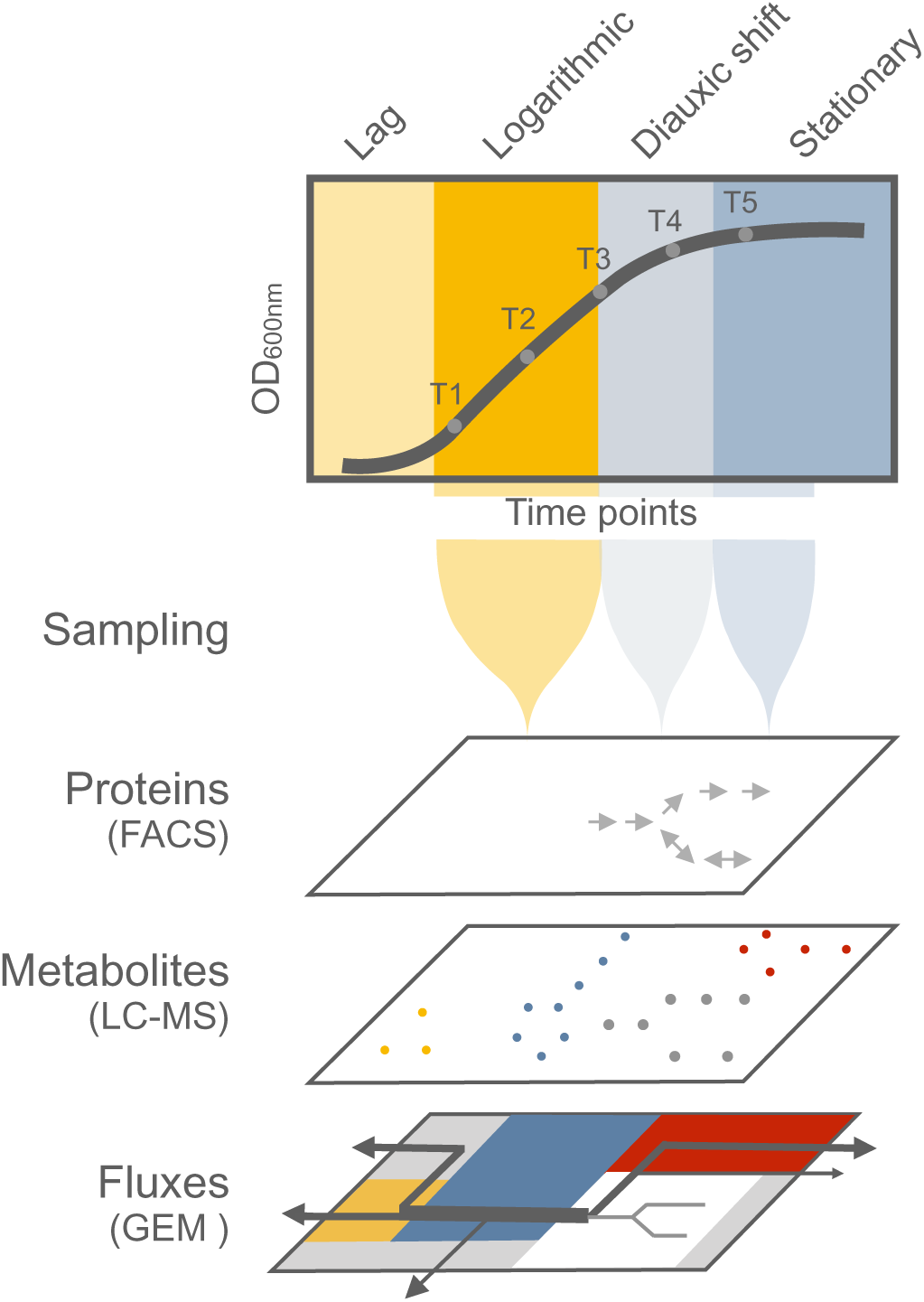

